# Anticipation of appetitive operant action induces sustained dopamine release in the nucleus accumbens

**DOI:** 10.1101/2022.07.26.501546

**Authors:** Jessica Goedhoop, Tara Arbab, Ingo Willuhn

**Author notes:** **Corresponding author email address** Ingo Willuhn –. **AUTHOR CONTRIBUTIONS** Jessica Goedhoop: Conceptualization; Data curation; Formal analysis; Investigation; Visualization; Methodology; Writing Tara Arbab: Conceptualization; Writing; Editing Ingo Willuhn: Conceptualization; Resources; Data curation; Supervision; Funding acquisition; Methodology; Writing; Editing; Project administration. **FUNDING** H2020 European Research Council (ERC): Ingo Willuhn, ERC-2014-STG 638013; Netherlands Organization for Scientific Research (NWO): Ingo Willuhn, VIDI 864.14.010,2015/06367/ALW; Netherlands Organization for Scientific Research (NWO): Ingo Willuhn, Gravitation program, BRAINSCAPES 024.004.012. The funders had no role in study design, data collection and interpretation, or the decision to submit the work for publication. The authors declare that no competing interests exist. **Code availability** The code used for this study is available from the corresponding author upon request.

## Abstract

The mesolimbic dopamine system is implicated in signaling reward-related information as well as in actions that generate rewarding outcomes. These implications are commonly investigated in either Pavlovian or operant reinforcement paradigms, where only the latter requires instrumental action. To parse contributions of reward- and action-related information to dopamine signals, we directly compared the two paradigms: rats underwent either Pavlovian or operant conditioning while dopamine release was measured in the nucleus accumbens, a brain region central for processing this information. Task conditions were identical with the exception of the operant-lever response requirement. Rats in both groups released the same quantity of dopamine at the onset of the reward-predictive cue. However, only the operant-conditioning group showed a subsequent, sustained plateau in dopamine concentration throughout the entire five-second cue presentation (preceding the required action). This dopamine “ramp” was unaffected by probabilistic reward delivery, occurred exclusively prior to operant actions, and was not related to task performance or task acquisition, as it persisted throughout the two-week daily behavioral training. Instead, the ramp flexibly increased in duration with longer cue presentation, seemingly modulating the initial cue-onset triggered dopamine release (i.e., the reward-prediction error (RPE) signal), as both signal amplitude and sustain diminished when reward timing was made more predictable. Thus, our findings suggest that RPE and action components of dopamine release can be differentiated temporally into phasic and ramping/sustained signals, respectively, where the latter depends on the former and presumably reflects the anticipation or incentivization of appetitive action, conceptually akin to motivation.

**Significance Statement:** It is unclear whether the components of dopamine signals that are related to reward-associated information and reward-driven approach behavior can be separated. Most studies investigating the dopamine system utilize either Pavlovian or operant conditioning, which both involve the delivery of reward and necessitate appetitive approach behavior. Thus, used exclusively, neither paradigm can disentangle the contributions of these components to dopamine release. However, by combining both paradigms in the same study, we find that anticipation of a reward-driven operant action induces a modulation of reward-prediction-associated dopamine release, producing so-called “dopamine ramps”. Therefore, our findings provide new insight into dopamine ramps, and suggest that dopamine signals integrate reward and appetitive action in a temporally distinguishable, yet dependent, manner.

## INTRODUCTION

Striatal dopamine plays a prominent role in motivated behavior and reward learning. More specifically, activity of the mesostriatal dopamine system is associated with movement (Lee et al., 2019; Howe and Dombeck 2016; Da Silva et al., 2018; Jin and Costa 2010; Schultz et al., 1983; Syed et al., 2016), motivational processes including the attribution of incentive salience (Berridge, 2007; Saunders et al., 2018; Salamone and Correa 2012; Flagel et al., 2011), reward value (Hamid et al., 2016), and the so-called temporal-difference reward-prediction error (RPE), central to reinforcement learning (Schultz et al., 1997; Hart et al., 2014; Bayer and Glimcher 2005; Sutton and Barto 1987). A long-standing question is how these aforementioned functions associated with motivation and learning are integrated into the release of dopamine from neuronal terminals in the striatum (Berke, 2018). Relatedly, it is not fully understood whether these functions depend on one another, or whether they govern striatal dopamine-release dynamics independently, distinct in *time* (i.e., RPEs are encoded at a different time point than the motivational drive to pursue a reward) or *space* (i.e., across different regional domains of the striatum).

Regarding *temporal* separation of dopamine-signal functions: theoretical conceptualizations distinguish slow changes (on the order of minutes) affecting “tonic” or ambient dopamine concentration, in contrast to much faster, “phasic” changes (Grace, 1991). Additionally, a number of studies report so-called ramping changes in dopamine release that are of intermediate speed, on the order of seconds (Lerner et al., 2021). Fast dopamine dynamics are proposed to serve a learning function (i.e., encode RPEs and other related value functions), whereas slow dynamics may be important for motivation (Niv, 2007). Consistent with their intermediate time scale, dopamine ramps are hypothesized to be influenced by a number of variables pertinent to either motivation or learning. These variables include reward expectation and proximity, RPE, state value, and uncertainty (Howe et al 2013; Gershman, 2014; Guru et al., 2020; Mikhael et al., 2022; Kim et al., 2020). Among these, state uncertainty is assumed to be of particular importance, as it is impactful and can theoretically explain the effect of the other variables (Starkweather et al., 2017, Mikhael et al., 2022; Kim et al., 2020).

Regarding *spatial* separation of dopamine-signal functions: differences in dopamine signaling across striatal regions have been reported extensively, demonstrating that representation of these functions varies by region (Willuhn et al., 2012; Willuhn et al., 2014; Lammel et al., 2011; DeJong et al., 2019; Menegas et al. 2017; Klanker et al. 2019; van Elzelingen et al 2022a; van Elzelingen et al 2022b). Although the resulting striatal dopamine landscape is not always consistent between studies, consensus is that midbrain dopamine neurons projecting to the nucleus accumbens core (part of the ventromedial striatum (VMS)), participate in both motivation and learning (van Elzelingen et al 2022b). Thus, the VMS is an optimal target to study the integration of dopamine signals encoding motivated actions required to earn rewards, as well as the RPEs associated with these rewards. In order to do so, we measured dopamine release in the VMS of rats undergoing either Pavlovian conditioning (PC) or operant conditioning (OC), using this direct comparison to parse the contributions of reward-related information (e.g.., reward-predictive cue) and motivated action to a dopamine signal.

Our results show that the onset of a reward-predicting cue induces an initial rise in dopamine release that is indistinguishable between PC and OC. However, we observe marked differences in the sustain of this dopamine release between PC and OC during the remainder cue presentation, where the only behavioral difference was that PC rats approached the food magazine and OC animals approached the location where the lever extended after cue offset. Together, our findings identify a ramping anticipation component of dopamine release (which is temporally separated from, yet dependent on, the RPE component) and provide new insight on how learning and motivation functions are integrated in VMS dopamine signals.

## RESULTS

### Behavior during Pavlovian and operant conditioning

Rats were trained on a variable intertrial-interval (VITI) of 60s using one of two experimental paradigms: either PC, in which a cue signals reward delivery, or OC, in which the same cue signals that a (required) lever press will produce reward delivery (Figure 1A). Rats undergoing PC (n = 12) received the maximum number of obtainable rewards (40) immediately from session 1 onwards, while OC rats (n = 13) on average earned the maximum number of rewards from approximately session 3 onwards (Figure 1B). This learning curve of the OC group was also reflected in the decreasing latency to lever-press after lever extension (main effect: χ^2^(14)=5268, p<0.0001), with sessions 3 through 14 having a significantly lower latency to lever-press compared to session 1 (1 vs. 3: p=0.0475; 1 vs. 4: p=0.0134; 1 vs. 5: p=0.0081; 1 vs. 6: p=0.0016; 1 vs. 7: p<0.0001; 1 vs. 8: p=0.0013; 1 vs. 9: p=0.0068; 1 vs. 10: p=0.0002; 1 vs. 11: p=0.0002; 1 vs. 12: p= 0.0002; 1 vs. 13: p=0.0017; 1 vs. 14: p<0.0001; Figure 1C). We restricted the following behavioral analysis to those sessions (1, 3, 6, 14) in which dopamine measurements were taken. The two groups differed in their appetitive-approach behavior during the 5s cue-light exposure: Over the course of conditioning, the PC group rapidly developed cue-induced approach behavior towards the reward magazine, illustrated by a high probability (Main effect of session: F(1.273,28.01)=0.4624, p=0.5481; Main effect of group: F(1,23)=1025, p<0.0001; Session x group interaction: F(3,66)=68.90, p<0.0001), much time spent (Main effect of session: F(2.021,44.45)=19.19, p<0.0001; Main effect of group: F(1,23)=742.6, p<0.0001; Session x group interaction: F(3,66)=67.94, p<0.0001), and low latency (Main effect of session: F(1.599,35.17)=1.249, p<0.0001; Main effect of group: F(1,23)=699.0, p<0.0001; Session x group interaction: F(3,66)=48.55, p<0.0001) to approach this section of the operant box, whereas the OC group did not (Figure 1D). *Post-hoc* analysis revealed that sessions 3, 6, and 14 differed significantly from session 1 in reward-magazine approach probability (PC: 1 vs. 3: p=0.0007; 1 vs. 6: p=0.0009; 1 vs. 14: 0.0012; OC: 1 vs. 3: 0.0002; 1 vs. 6: p=0.0001; 1 vs. 14: 0.0004), time spent (PC: 1 vs. 3: p<0.0001 ; 1 vs. 6: p<0.0001; 1 vs. 14: p=0.0001; OC: 1 vs. 3: p<0.0001; 1 vs. 6: p<0.0001; 1 vs. 14: 0.0002), and latency (PC: 1 vs. 3: p=0.0017; 1 vs. 6: p=0.0017; 1 vs. 14: p=0.0039; OC: 1 vs. 3: p=0.0001; 1 vs. 6: p<0.0001; 1 vs. 14: p=0.0004); sessions 3, 6, and 14 did not, however, differ significantly from each other, indicating that from session 3 onwards, conditioned approach behavior was stable and no additional learning occurred after this time point. In contrast to the PC group, the OC group rapidly developed cue-induced approach behavior towards the lever below the cue light, which they were trained to press after the cue light turned off in order to obtain a food pellet. This, too, was illustrated by a high probability (Main effect of session: F(1.468,31.81)=14.23, p=0.0002; Main effect of group: F(1,23)=303.1, p<0.0001; Session x group interaction: F(3,65)=20.50, p<0.0001), much time spent (Main effect of session: F(2.183,47.30)=26.93, p<0.0001; Main effect of group: F(1,23)=174.7, p<0.0001; Session x group interaction: F(3,65)=30.88, p<0.0001), and a low latency (Main effect of session: F(1.899,41.15)=16.10, p<0.0001; Main effect of group: F(1,23)=345.8, p<0.0001; Session x group interaction: F(3,65)=19.33, p<0.0001) to approach the lever during the 5s cue exposure (Figure 1E). Again, *post-hoc* analysis revealed that sessions 3, 6 and 14 differed significantly from session 1 in lever-approach probability (1 vs. 3: p<0.0001 ; 1 vs. 6: p<0.0001 ; 1 vs. 14: p=0.0003), time spent (1 vs. 3: p<0.0001 ; 1 vs. 6: p<0.0001; 1 vs. 14: p=0.0002), and latency (1 vs. 3: p=0.0003 ; 1 vs. 6: p<0.0001 ; 1 vs. 14: p=0.0004); and sessions 3, 6 and 14 did not differ significantly from each other, indicating that in the OC group as well, from session 3 onwards, conditioned approach behavior was stable and no additional learning occurred after this time point. The number of PC-group approaches to the lever or cue light did not differ significantly between sessions. Thus, although the two groups approached different areas of the operant box, both groups learned this appetitive approach behavior at the same rate (i.e., by day 3); this is also demonstrated by the latencies of their respective cue-induced approaches, which did not differ between the two groups (Main effect of group: F(1,23)=2.140, p=0.1571; Figure 1F). Conceptually, this latency might also reflect approach vigor, which implies that vigor also did not differ between the two groups. To further investigate this last point, we compared the locomotion speed of the groups during the 5s cue-light exposure (Figure 1G). Locomotion speed between PC and OC groups differed significantly in sessions 1 (U=20, p=0.0041) and 3 (U=35, p=0.0188), but not sessions 6 (U=61, p=0.3760) and 14 (U=35, p=0.1072). Additionally, the groups did not differ in their distance to the reward magazine during the 5s-cue exposure in session 1 (U=49, p=0.3434), however, there was a significant difference in subsequent sessions (3: U=0, p<0.0001; 6: U=0, p<0.0001; 14: U=0, p<0.0001; data not shown), when the PC group had learned to approach the magazine and the OC group had learned to approach the lever instead. Taken together, we conclude that approach vigor overall (measured as approach latency and speed), did not differ between groups. Once animals had acquired their respective approach behavior, they spent most of the time during cue exposure near the approached object, and moved away from the food magazine after reward consumption.

**Figure 1.**
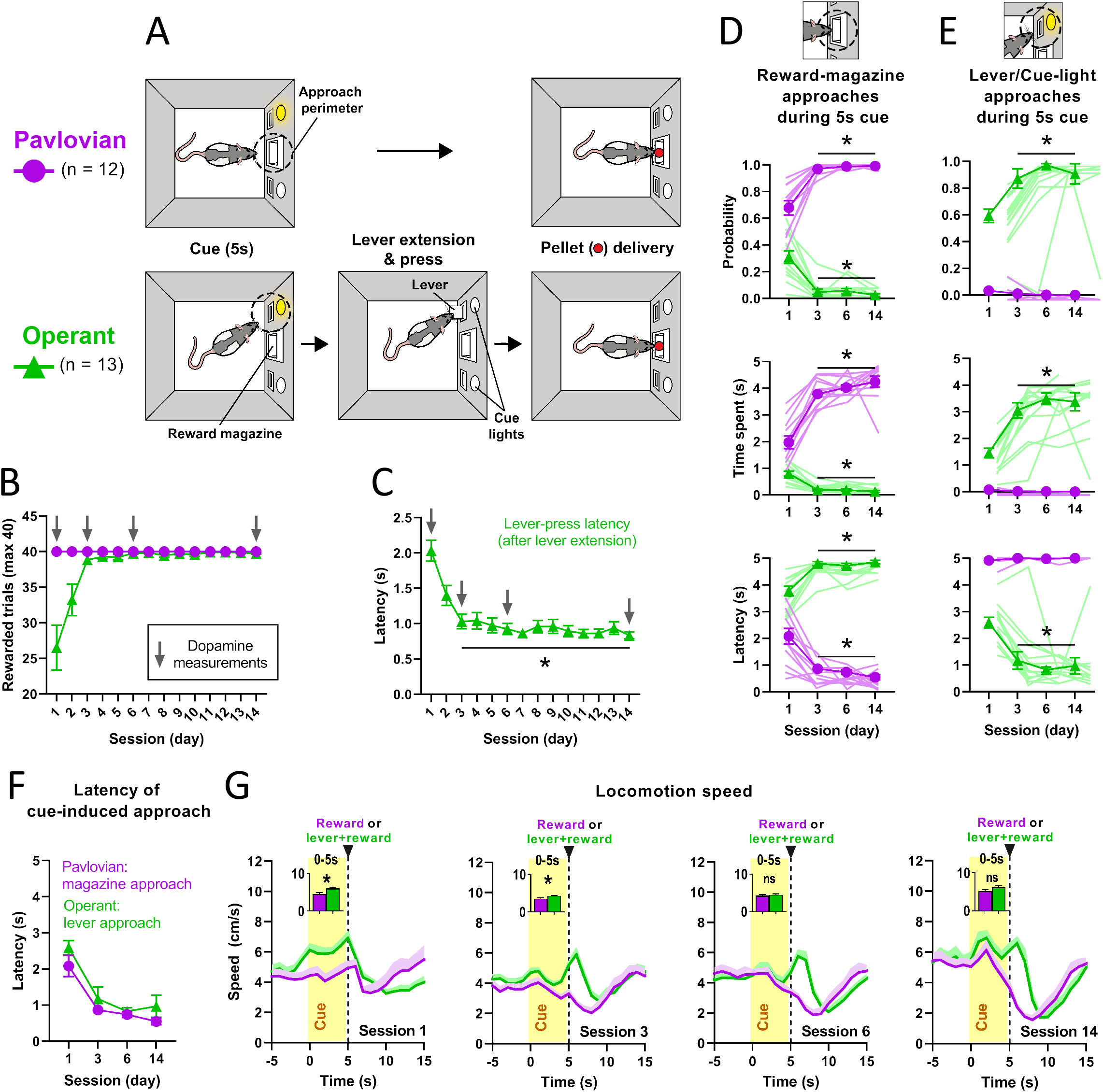
Behavior during Pavlovian and operant conditioning. **A** Schematic of the PC and operant task. For the PC group (n=12, purple circles/traces throughout all figures) a 5s cue-light exposure was followed by the immediate delivery of a single food pellet. For the OC group (n=13, green triangles/traces throughout all figures) a 5s cue-light exposure was followed by the extension of a lever, which needed to be pressed within 5s in order for the delivery of a single food pellet. **B** Average number of rewarded trials (maximum of 40 per session) over the course of conditioning sessions. Arrows mark FSCV-recorded sessions. **C** Average latency to lever press after lever extension decreases over the course of conditioning for the OC group. **D** Group differences in cue-induced reward-magazine approach are reflected by probability, time spent, and latency during the 5s cue-light. Reward-magazine approach stabilized from session 3 onwards. **E** Group differences in cue-induced lever/cue-light approach behavior are reflected by probability, time spent, and latency during the 5s cue-light exposure (Statistics available at https://osf.io/jhz7x/). **F** Average latency of the respective cue-induced approach (PC group: towards reward magazine; OC group: towards lever) did not differ between the two groups. **G** Average locomotion speed across recording sessions. Bar graph insets depict speed restricted to the 5s cue-light exposure, which differed significantly between groups during sessions 1 and 3, but not 6 and 14. All data are mean±SEM. Single-animal data are represented in lighter-shaded lines in D&E. * = p ≤ 0.05.

### Dopamine release in the VMS during Pavlovian and operant conditioning

Extracellular dopamine fluctuations were measured during PC and OC sessions using fast-scan cyclic voltammetry (FSCV), with chronic electrodes targeting the VMS (Figure 2A). Both groups released dopamine in response to cue presentation as exemplified by the representative color plots in Figure 2B. In addition, both groups stably released dopamine in response to the delivery of unpredicted food pellets, given as a control for electrode viability prior to the start of each session; there was no difference between groups (Main effect of time: F(2.618,56.73)=0.8803, p=0.4448; Main effect of group: F(1,23)=1.983, p=0.1724; Figure 2C). Average dopamine release during the 5s-cue exposure (time epoch: 0-2.5s and 2.5-5s, with baseline set before cue-light illumination) and reward delivery (time epoch: 5-7.5s, with baseline set before pellet delivery) remained relatively stable over time (Figure 2D), where dopamine release did not differ between groups at initial cue onset (0-2.5s) or after reward delivery, but did differ during cue exposure (2.5-5s; see below). More specifically, while dopamine induced by cue exposure (2.5-5s) did not differ between groups in session 1 (t(21)=0.0612, p=0.9518), it differed significantly in subsequent sessions (3: t(23)=2.686, p=0.0132; 6: t(23)=2.188, p=0.0391; 14: t(20)=2.147, p=0.0443; Figures 2D, bottom, and 2E). The dynamics are characterized respectively by an initial rise (which is known to track changes in reward value and encodes an RPE; Figure 2F) followed by an immediate drop towards baseline dopamine concentration for the PC group, while cue-induced dopamine in the OC group was sustained throughout the 5s-cue exposure (which the PC group showed only in session 1). Reward-induced changes in dopamine due to reward delivery did not differ between groups throughout sessions (1: U=47, p=0.2839; 3: U=59, p=0.3203; 6: U=51, p=0.1519; 14: U=43, p=0.2829; Figure 2D, top). In addition in neither of the groups, cue-induced dopamine release (time epoch: 2.5-5s) was correlated to locomotion speed, approach probability, approach time spent, approach latency nor distance to the food magazine in any of the conditioning sessions (Figure 2G).

**Figure 2.**
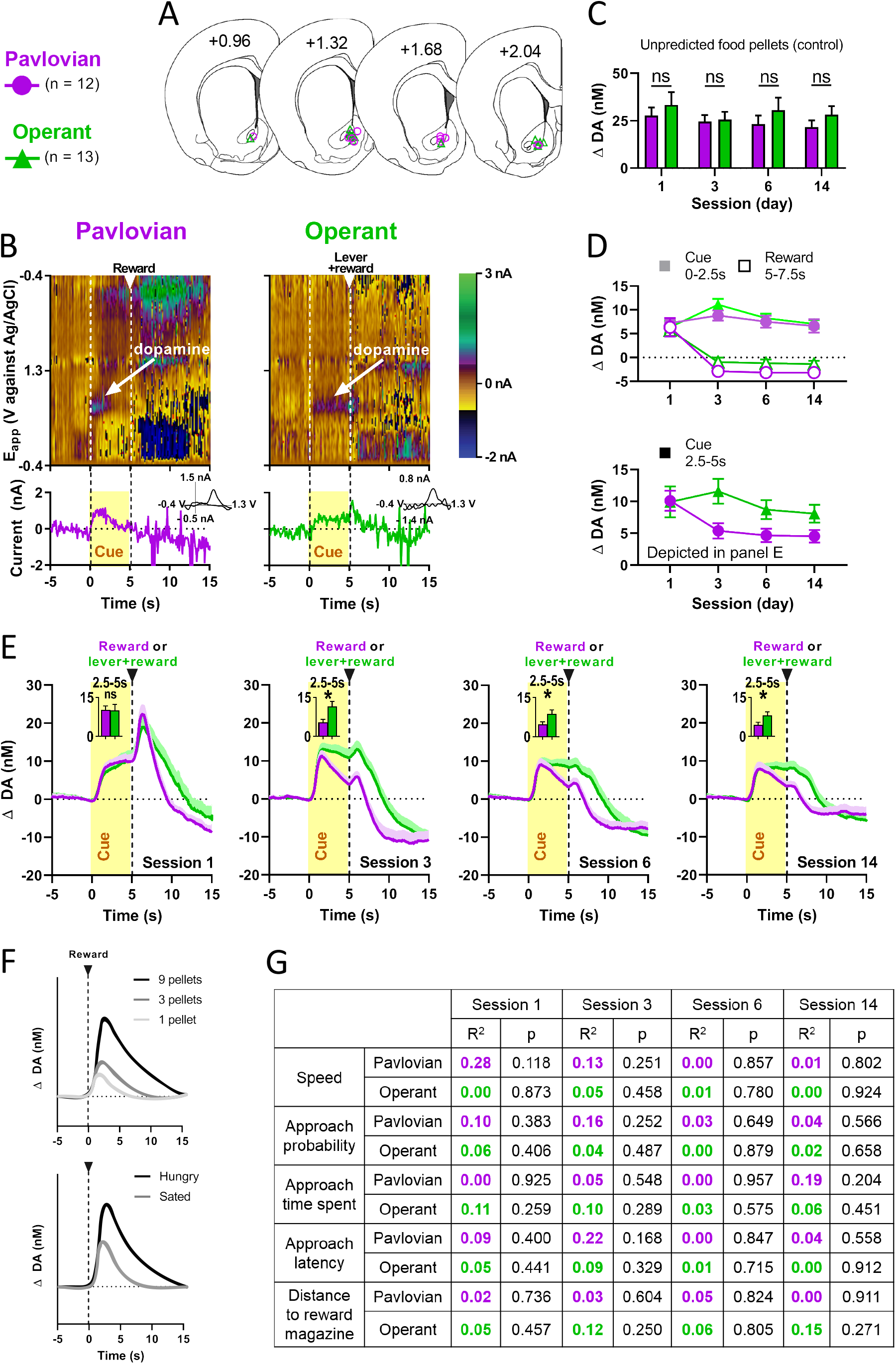
Dopamine release in the VMS during Pavlovian and operant conditioning. **A** Histological verification of electrode placement in the VMS. Purple circles represent rats from the PC group (n=12) and green triangles represent rats from the OC group (n=13). **B** Example single-trial pseudo-color plots (top) and dopamine traces (bottom) and cyclic voltammograms (bottom insets) of dopamine recordings in PC (left) and OC (right) groups. Represented are 5s before cue-light exposure, 5s of cue light exposure, lever-press and/or food pellet delivery, and 10s after the cue-light exposure. **C** Average peak values of dopamine release in response to unpredicted food pellets. No significant differences were found between PC and OC groups during any of the sessions. **D** Top: Average cue- (time epoch: 0-2.5s, baseline set before start cue-light exposure) and reward- (time epoch: 5-7.5 s, baseline set before food pellet delivery) induced dopamine release. Bottom: Average cue-induced (time epoch: 2.5-5s, baseline set before start cue-light exposure) dopamine release. **E** Average dopamine concentration across seconds (per session). We observed sustained dopamine release during the 5s cue-light in the OC group in all four sessions, in contrast to the PC group, which only showed sustained release during session 1 (time epoch: 2.5-5s). **F** The phasic dopamine peak immediately following a stimulus is known to track changes in reward value and encodes an RPE, as dopamine scales with the unexpected delivery of rewards of different value (top: number of food pellets; bottom: hunger state). Modified from van Elzelingen et al (2022b). **G** There were no correlations between average cue-induced dopamine release (time epoch: 2.5-5s) and locomotion speed, approach probability, approach time spent, approach latency or distance from reward magazine. All data are mean±SEM. Dopamine data are baseline subtracted. * = p ≤ 0.05.

In order to better understand the groups’ different dopamine dynamics during the 5s-cue exposure, several additional training and recording sessions were performed in which we varied task-relevant parameters. For the first experiment, we set out to further investigate how the anticipation to perform an action (the lever press) affects dopamine. We prolonged the duration of anticipation to lever press by increasing the cue-light duration to 10s, which resulted in a significant difference in average dopamine release between groups (time epoch: 5-10s, with baseline set before cue illumination; t(17)=1.759, p=0.0483), with the OC group (n=10) showing sustained dopamine release for the entire cue period in contrast to the PC group (n=9, Figure 3A).

**Figure 3.**
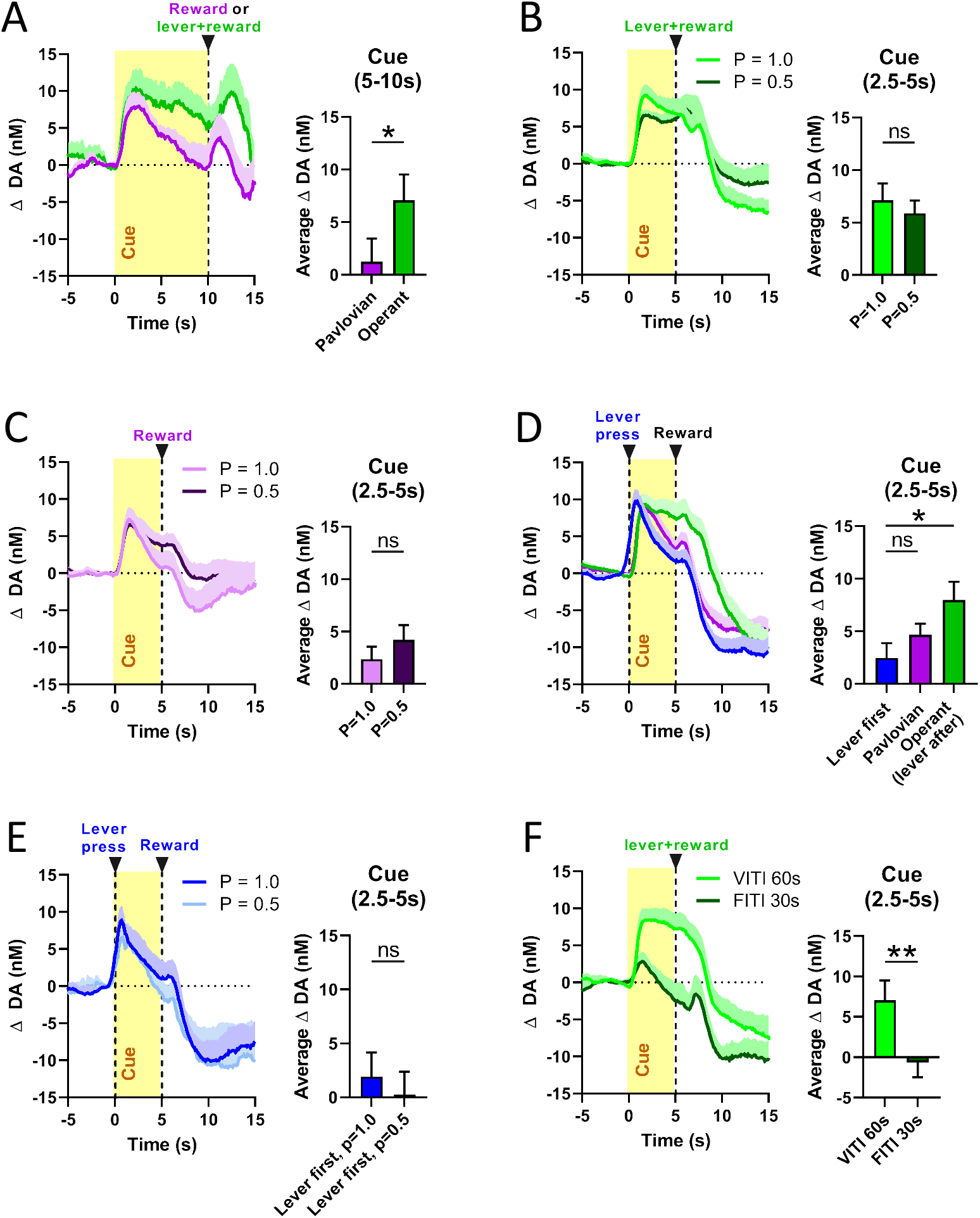
Sustained dopamine release during cue presentation reflects the anticipation of appetitive action. **A** Increasing the duration of cue-light exposure from 5s to 10s prolonged sustained dopamine release (time epoch: 5-10s) in the OC group (n=10) compared with the PC group (n=9). **B** Lowering the probability of food-pellet delivery from P=1.0 to P=0.5 did not change sustained cue-induced dopamine release in the OC group (n=9). **C** Lowering the probability of food-pellet delivery from P=1.0 to P=0.5 did not induce sustained cue-induced dopamine release in the PC group (n=8). **D** Requiring a lever press prior to 5s cue-light exposure and food-pellet delivery (lever-first group, n=8) eliminated sustained dopamine release, as cue-induced dopamine release (time epoch: 2.5-5s) did not differ between the lever-first and PC groups, but differed significantly between the lever-first and OC groups. **E** Lowering the probability of food-pellet delivery from P=1.0 to P=0.5 when a lever press is required prior to cue and pellet delivery did not affect average cue-induced dopamine release (time epoch: 2.5-5s; n=4). **F** Changing the ITI from a VITI-60s to a FI-30s for the OC group resulted in a significant decrease in average dopamine release (n=7) and no more sustained dopamine release. All data are baseline subtracted, mean±SEM. Left, average dopamine-release traces. Right, bar graphs depicting dopamine-release restricted to the cue-exposure. * = p ≤ 0.05, ** = p ≤ 0.01.

Further, we hypothesized that OC-group rats may experience greater uncertainty about receiving the food pellet, since reward delivery was contingent on lever pressing (despite fulfilling this lever-press requirement correctly in almost all trials from session 3 onwards), and that this uncertainty might induce more sustained dopamine release during the 5s-cue exposure compared to the PC group (which had no operant requirement). This idea is supported by the fact that, in the PC group, cue-induced dopamine release was similarly sustained in session 1, in which the rats were still learning the contingency between cue and reward and, thus, experienced greater uncertainty about receiving reward. We increased uncertainty by decreasing the probability of food pellet delivery from P=1.0 to P=0.5. However, this uncertainty did not significantly alter cue-induced dopamine release in OC (Z=-1.125, p=0.2604; n=9; Figure 3B) or PC (Z=-1.540, p=0.1235; n=8; Figure 3C) groups.

In another experiment, we eliminated the element of anticipation to lever press from the 5s cue-light period: We switched the order of appearance of lever and cue light in the OC group, so that the VITI-60s was followed by immediate extension of the lever into the operant box, and a lever press (within 5s of lever extension) resulted in a 5s cue-light illumination, after which the light turned off and a reward was delivered. As a consequence, rats (‘lever-first’ group, n=8) did not show sustained dopamine release during the cue light, and their dopamine dynamics resembled that of the PC group (Figure 3D); the latter observation is supported by the fact that mean dopamine concentration during cue exposure did not differ significantly between lever-first and PC groups conditioned for a similar number of days (Main effect: F(2,30)=3.310, p =0.0502; *post-hoc* testing: p=0.5101), yet differed significantly from rats that underwent regular OC (lever-press required after the cue light turned off; p=0.0338). In a subset of these “lever-first” rats, we decreased the probability of food pellet delivery from P=1 to P=0.5 (n=4), which again did not significantly alter cue-induced dopamine release (t(3)=0.8821, p=0.2213; Figure 3E).

All previously described experiments made use of a VITI of 60s (range: 30-90s), thus, rats were not able to predict when in the session the next trial would start (i.e., when the cue light is turned on). In a final experiment, we tested whether such “session uncertainty” was a prerequisite for the observed effects by using a highly predictable, fixed ITI of 30s, which eliminated almost all cue-induced dopamine (Figure 3F). No more sustained dopamine release was observed in the OC group, and the average dopamine concentration was significantly lower compared to VITI-60s conditions (n=7, t(6)=3.418, p=0.0071).

## DISCUSSION

To investigate how the mesolimbic dopamine system integrates functions related to reward and actions associated with the pursuit of reward, we measured VMS-dopamine release in rats undergoing either PC or OC, in which a visual cue signaled either the upcoming delivery of a food pellet, or the opportunity to execute an action to obtain this reward, respectively. Initial dopamine-release amplitude to the cue was similar between groups, but in OC, we observed a sustained elevation of dopamine concentration subsequently (throughout cue presentation and prior to lever press) compared to PC. This dopamine sustain or ramp was observed reliably and consistently throughout systematic manipulation of experimental parameters and behavioral training, and, thus, we interpret it as associated with the anticipation or preparation to execute the (learned) operant action.

Our parallel PC/OC study design utilized two paradigms that are nearly identical and differ only by a brief instrumental action in OC (operant-lever response requirement), which takes place in close physical proximity to the food magazine (following the offset of the 5s-cue presentation). The onset of the cue induces immediate appetitive approach-behavior in both groups of rats, distinguishable only by its target location (OC rats approached the lever site and PC rats the food magazine). During the 5s-cue epoch both groups remain near their respective target location, whereas only after cue offset behavior differed momentarily, as PC rats consumed the food immediately and OC rats performed a brief, single lever-press just prior to food consumption. Both groups acquired the approach behavior with a similar time course, where behavior was already stable after 2-3 of the 14 training sessions. Thus, because overall behavior during cue presentation did not differ between groups, we can exclude a number of explanations for the observed differences in sustained dopamine release. Such differences cannot be attributed to learning, as group behavior did not differ in learning rate, and dopamine signaling also developed with a similar time course in both groups (2-3 sessions). Furthermore, post-learning behavior (extended behavioral training (14 sessions)) during cue presentation was similar in both groups, indicating that approach behavior, task performance (Fig 1d-g; Fig 2f; including response vigor), and general speed of movement were not sources for differential dopamine signaling. Furthermore, differential dopamine dynamics can not be explained by varying electrode sensitivity, which was stable across training and groups. Taken together, because of the aforementioned similarity of OC and PC behavior (prior to lever press), our findings suggest that sustained OC dopamine is related to behavior that has not yet occured: the upcoming lever press after cue offset.

To further interrogate the nature of sustained dopamine release, we systematically varied behavioral-paradigm parameters. First, we extended the duration of the predictive cue from 5s to 10s, which extended the sustained release until lever press and, thus, demonstrated that this dopamine ramp is flexible in duration and is associated with a “state” that precedes action initiation. Such a state may for example be linked to the “readiness” to perform an action and may bridge the time between action anticipation and its execution. Next, since previous work demonstrated that the dopamine system is sensitive to reward uncertainty (Kobayashi and Schultz, 2008; Fiorillo et al., 2003; de Lafuente and Romo, 2011; Starkweather et al., 2017; Lak et al., 2017), we rendered food delivery probabilistic (only 50% of trials rewarded, versus 100% in previous experiments). In neither PC or OC did such uncertainty lead to a significant change in sustained dopamine. In contrast, the dopamine ramp disappeared after we moved the lever-press requirement forward in time, from cue offset to cue onset; thus, sustained dopamine is only observed with an operant requirement that *follows* a period of action anticipation. In this “lever-first” situation, increasing uncertainty by rewarding only 50% of trials did not reinstate sustained dopamine, further underlining insensitivity of this ramp phenomenon to uncertainty. Thus, dopamine ramps were not a simple product of unexpected reward prediction (variable ITIs prevent the animals from predicting time of trial onset) and operant requirement, but instead of a specific sequence of the two: reward-predicting cue followed by lever press. Moreover, this dopamine ramp instantiates a modulation or extension of the initial phasic component of dopamine release (known to constitute an RPE signal (van Elzelingen et al., 2022b; Hart et al., 2014; Figure 2F), as both amplitude and sustain of dopamine ramps diminish drastically when the timing of reward delivery is (more) temporally predictable (fixed ITI). Interestingly, the initial dopamine peak had the same amplitude in both groups, indicating no value discounting (e.g., due to the required effort), as well as that the ramp did not consist of a redistribution of RPE-related dopamine signaling. Therefore, taken together, the two core elements necessary for dopamine ramps constitute a positive RPE followed by an operant-action anticipation necessary to earn this reward, neither of which were sufficient to produce a dopamine ramp on their own.

Ruling out movement as a direct source for changes in dopamine signaling seems, at first glance, inconsistent with several previous studies. However, most studies that tie the dopamine system to initiation of specific movement, vigor, or velocity were executed on the single-cell level (Da SIlva et al., 2018; Dodson et al., 2016; Howe and Dombeck 2016; Barter et al., 2015; Hughes et al., 2020; Engelhard et al., 2019; Coddington and Dudman 2018; Jin and Costa 2010; [Puryear et al., 2010; Wang and Tsien, 2011), whereas our study evaluates “bulk” signal from dopamine-neuron terminals, reflecting dopamine released from many neurons, which likely dilutes movement-specific activity. Indeed, bulk-signal studies report more general associations with movement (Syed et al., 2016; Lee et al., 2019; Flagel et al., 2011). Notably, the dopamine system is set up anatomically to broadcast its signals to striatal targets via bulk or population signals, as dopamine is released from a large number of extra-synaptic terminals, resulting in a diffusion-based signal that is perpetuated by volume transmission (Rice & Cragg, 2008). Furthermore, terminal release can be modified independently of cell-body activity, which may contribute to a discrepancy in findings between sampling dopamine cell bodies and terminals (Threlfell et al., 2012). Relatedly, our results indicate a fusion or dependency of RPE and action-anticipation signals, consistent with a report that suggests a dependence of “dopaminergic” reward processing on movement related to reward pursuit (Syed et al., 2016). However, in our case this dependence is inverted, i.e., the action-anticipation signal is dependent on the RPE signal. Together, our results therefore suggest that unlike single-dopamine neuron-activity in the midbrain, bulk terminal dopamine release does not encode movement control. However, our findings nonetheless underline the link between dopamine RPE and movement, albeit with a not yet executed, anticipated movement.

So-called dopamine ramps are reported to occur over a timescale of seconds in both the dopaminergic midbrain as well as the ventral striatum, often when dopamine neuron-activity (including dopamine release) was sensed as bulk activity (Hamid et al., 2016; Howe et al., 2013; Collins et al., 2016; Kim et al., 2020; Guru et al., 2020). Such ramps are often associated with the gradual approach towards reward, or sensory feedback via stimuli that update about the prediction of impending reward (Kim et al., 2020; Mikhael et al., 2022) and, as in our hands, are more readily observed in OC compared to PC (Hamid et al., 2021; Guru et al., 2020; Song and Lee, 2020). However, in our experiment, the “amount” of sensory feedback (cue) and the animals’ distance to reward (food magazine) remain stable (until cue offset) in both OC and PC animals. Thus, it is possible that multiple sequential cues until reward delivery would have induced a ramp in both OC and PC animals, but in the absence of such scenarios, only OC animals show a ramp. Another inconsistency with previous reports is that we find remarkable stability of dopamine ramps across 14 days of behavioral training, whereas others have suggested and reported that ramps fade with extended training, when task performance becomes asymptotic (Collins et al., 2016; Guru et al., 2020; Song and Lee, 2020), potentially linking credit back to the rewarded action to guide reward learning Hamid et al., 2016; Howe et al., 2013; Collins et al., 2016). A function that would require such stable dopamine ramps is the encoding of reward expectation (Howe et al., 2013; Mohebi et al., 2019). However, in our paradigm, OC and PC rats had the same reward expectation, but only OC animals exhibited ramps. Together, this favors another explanation, the anticipation of performing a rewarded action: dopamine ramps could support the motivation to perform operant actions for “distal” rewards. Indeed, many previous findings tie the mesolimbic dopamine system to motivation (Salamone et al., 2007; Nicola, 2010; Berridge, 2007; Hughes et al., 2020; Panigrahi et al., 2015) and, moreover, some studies support the idea that dopamine ramps are implicated in motivation (Hamid et al., 2016; Wassum et al., 2012; Howe et al., 2013; Collins et al., 2016; Mohebi et al., 2019).

In summary, our findings suggest that sustained dopamine release during presentation of a reward-predicting cue can be driven by action anticipation, by means of modulating the ramp-preceding dopamine RPE signal (which is a precondition to the dependent ramp). These findings shine light on how the mesolimbic dopamine system integrates reward- and action-related functions associated with the pursuit of rewarding outcomes in a temporally distinguishable manner, and provide new insight into the nature and function of so-called dopamine ramps, which may embody an intermediate between learning and action, conceptually related to the motivation to generate a reward-achieving action.

## MATERIALS AND METHODS

### Animals

Adult male Long Evans rats (300-400g) were individually housed and kept on a reversed 12h day-night cycle (light on from 20:00 to 08:00) with controlled temperature and humidity. A total of 37 rats underwent surgery and were randomly assigned to either PC or OC experimental groups. Due to a non-functional or misplaced FSCV electrode, 12 rats were excluded from this study and the final group sizes were n = 12 for the PC experiment and n = 13 for the OC experiment. The rats were food restricted to 85% of their free-feeding body weight, and water was provided *ad-libitum*. The rats underwent one training session per day, consistently at the same time. All animal procedures were in accordance with the Dutch and European laws and approved by the Animal Experimentation Committee of the Royal Netherlands Academy of Arts and Sciences.

### Stereotaxic surgery

Rats were anesthetized using isoflurane (1-3%) and placed into the stereotactic frame. Body temperature was maintained using an isothermal pad. The analgesic Metacam (0.2mg meloxicam/100 g) was delivered using a subcutaneous injection and the shaved scalp was disinfected using ethanol (70%). An incision of the scalp, which was treated with lidocaine (100mg/ml), exposed the cranium at the midline. A craniotomy was drilled and the dura mater was cleared in order to unilaterally target the nucleus-accumbens core of the VMS (1.2mm AP, 1.5mm ML and - 7.1mm DV) with a chronically-implanted carbon-fiber electrode (Clark *et al*., 2010) made in-house. An Ag/AgCl reference electrode was positioned in a separate part of the forebrain. The electrodes were secured with cranioplastic cement which was anchored to the skull by surgical screws. Rats received a subcutaneous injection of 2ml saline following surgery, and were placed in a temperature-controlled cabinet to be monitored for an hour. Rats were given 1-2 weeks post-surgery to recover before food restriction, behavioral training, and recordings started.

### Magazine- and FR1 training

All experiments were carried out in modified operant boxes (32 × 30 × 29cm, Med Associates Inc.), equipped with a food magazine (connected to an automated food-pellet dispenser) flanked by two retractable levers with cue lights above these levers, a house light, a white-noise generator and metal grid floors (Med Associates Inc.). Each operant box was surveilled by a video camera. The boxes were housed in metal Faraday cages that were insulated with sound-absorbing polyurethane foam. In order to habituate the rats to these operant boxes prior to conditioning and to teach the rats that they could obtain food pellet rewards (Dustless precision pellets, 45mg, Bio-Serv), the PC group received two magazine training sessions and the OC group received one magazine training session followed by a fixed ratio 1 (FR1) training session. During all training and recording sessions described in this paper, the house light was illuminated and white-noise was played at an intensity of 65dB to mask background noises. During magazine training sessions, 45 pellets were delivered on a variable intertrial-interval (VITI) of 90s (range 70 - 110 s). During the FR1 training session, the active lever (on the left side of the reward magazine) was extended into the operant box at the start of the session and each lever press resulted in the delivery of one food pellet. In this session, a maximum of 45 food pellets could be earned.

### Pavlovian and operant conditioning

For PC and OC, the rats were placed into the operant boxes and at the start of the house light was illuminated, the background whitenoise was turned on, and a VITI of 60s (range 30-90 s) was initiated. Following the ITI, a cue light was illuminated for a duration of 5s (Figure 1A). For the PC group, a food pellet was delivered into the reward magazine directly after the cue light turned off, after which the next VITI-60s started. For the OC group, the cue light turning off was followed by extension of the lever below the cue light into the operant box. The lever was retracted after one lever press (FR1), which immediately resulted in the delivery of one food-pellet reward into the food magazine, after which the next VITI-60s started. If there was no lever press within 5s after lever extension, the lever was retracted, no reward was delivered, and the next trial was started. The training sessions consisted of 40 trials; after the termination of the session, rats were returned to their homecage.

Rats underwent 22 consecutive daily training sessions, of which FSCV recordings took place on days 1, 3, 6, 14 and 22. On day 22 the amount of trials was increased to 80 and the regular trials (described above) were semi-randomly intermixed with trials in which an increased reward size was delivered (data not shown in this paper) or trials in which the cue-light illumination was prolonged to a duration of 10 s. Afterwards, a subset of the animals (PC group: n = 8, OC group: n = 9) was retrained on the regular training schedule for 3 days; on the fourth day, FSCV recordings took place throughout a session consisting of 20 regular trials, followed by 60 trials in which the probability of reward delivery was decreased to P=0.5. A subset of the rats from the OC group (n = 8) subsequently received 7 additional days of training with each session consisting of 40 trials in which the contingency was changed: During these sessions the VITI-60s was followed by immediate extension of the lever, and a lever press (within 5s of lever-extension) resulted in 5s cue-light illumination, after which the light turned off and a food pellet was always delivered. FSCV recordings took place on the seventh day of training on this schedule. During this recording session, a subset of the rats (n = 4) received an additional 40 trials in which the probability of reward delivery was decreased to P=0.5. For six PC and seven OC rats, training concluded with seven days of sessions with a fixed ITI of 30s, instead of the VITI of 60s during regular training. These sessions consisted of 40 trials and FSCV recordings took place on day 7.

### FSCV measurements and analysis

Fast-scan cyclic voltammetry (FSCV) was employed to detect subsecond changes in extracellular dopamine concentration, as was described previously (Willuhn et al., 2014). Chronically-implanted carbon-fiber micro-electrodes were implanted, and connected to a head-mounted voltammetric amplifier which was interfaced with a PC-driven data-acquisition and analysis system (National Instruments) through an electrical commutator (Crist) mounted above the test chamber. Voltammetric scans were repeated every 100 ms (10Hz). The electric potential of the carbon-fiber electrode was linearly ramped from -0.4V versus Ag/AgCl to +1.3V (anodic sweep) and back (cathodic sweep) at 400V/s (8.5ms total scan time) during each voltammetric scan, and held at -0.4V between scans. If present at the surface of the electrode, dopamine is oxidized during the anodic sweep resulting in the formation of dopamine-o-quinine (peak reaction detected around +0.7V), which is thereafter during the cathodic sweep reduced back to dopamine (peak reaction detected around -0.3V). The ensuing flux of electrons is measured as current and is directly proportional to the number of molecules that undergo electrolysis. The background-subtracted, time-resolved current obtained from each scan provides a chemical signature characteristic of the analyte, allowing resolution of dopamine from other substances (Phillips & Wightman, 2003). To isolate dopamine from the voltammetric signal, chemometric analysis with a standard training set was used (Clark et al., 2010). A moving 10-point median filter was used to smooth all data and baseline subtraction was performed on a trial-by-trial basis prior to analysis of average concentration. At the start of each FSCV recording session, two unexpected deliveries of a single food pellet (spaced apart by two minutes) confirmed electrode viability to detect dopamine. Animals were excluded from analysis when: 1) No dopamine was detected in response to the unexpected pellets before start of the behavioral session, or 2) FSCV recording amplitude background noise was larger than 1nA in amplitude.

### Behavioral analysis

The delivered rewards and (latency of) lever presses were registered via an automated procedure using MedPC (Med Associates Inc.). DeepLabCut software (Mathis et al., 2018) was used to track the position of the rats in the operant box on video data acquired during FSCV measurements. The tracking data was analyzed using MATLAB (The Mathworks, Inc. Version 2019a) to determine the distance of the rats to the reward magazine and lever (measured from the headcap), and the speed of movement (cm / s, measured from the middle of the back). To determine the probability, time spent, and latency of approaching the reward magazine or lever during the cue epoch, approaches were defined by the proximity of the rat to the reward magazine or lever of less than 5cm for at least 1s. The probability to approach was calculated using (number of cue exposures with at least 1s of approach in a session/total cue exposures in a session). Time spent approaching was calculated by averaging the time the rats spent approaching during the cue exposures in a session. The latency to approach was determined by averaging the latencies of the first approach (of at least 1s) during the cue exposures in a session. The average locomotion speed was determined during the cue epoch.

### Histological verification of recording sites

After completion of the experiments, rats were deeply anesthetized using a lethal dose of pentobarbital. Recording sites were marked with an electrolytic lesion before transcardial perfusion with saline, followed by 4% paraformaldehyde (PFA). Subsequently, the brains were removed and post-fixated in 4%-PFA for 24 hours before they were placed in 30%-sucrose for cryoprotection. After saturation, the brains were rapidly frozen in an isopentane bath, sliced on a cryostat (50µm coronal sections, -20°C), and stained with cresyl violet in order to increase the visibility of the electrode-induced lesions and anatomical structures.

### Statistical analysis

For all analyses only rewarded trials were included. Behavioral data was analyzed using one- or two-way repeated measures ANOVAs, unpaired T-tests, or their nonparametric equivalents when appropriate. *Post-hoc* analyses were conducted when necessary and p-values were adjusted when multiple comparisons were made. Average extracellular dopamine concentrations during the last half of the cue epoch (2.5s in case of a 5s cue and 5s in case of a 10s cue, with baseline set before cue-light illumination) and reward epoch (5 - 7.5s after cue onset, with baseline set before reward delivery) were compared between groups using unpaired T-tests or a one-way ANOVAs, or their nonparametric equivalents when appropriate. For the recordings in which the probability of reward delivery decreased within the session (P=1 to P=0.5), we used the average of the last 20 P=0.5 trials, excluding the trials in which the animals were learning the new contingency. In the case of within-subject comparisons, a paired T-test or its nonparametric equivalent was used. Regression analyses were performed to test for correlations between the average locomotion speed, approach probability, approach time spent, approach latency, average distance from the reward magazine, and average dopamine concentrations during the cue epoch.

## ACKNOWLEDGEMENTS

We thank Ralph Hamelink and Nicole Yee for their technical support. This research was funded by the European Research Council (ERC; ERC-2014-STG 638013) and the Netherlands Organization for Scientific Research (NWO; 864.14.010, 2015/06367/ALW; BRAINSCAPES 024.004.012). A CC BY or equivalent license is applied to the author’s accepted manuscript (AAM) arising from this submission, in accordance with the grant’s open access conditions.

